# Limited marginal utility of deep sequencing for HIV drug resistance testing in the age of integrase inhibitors

**DOI:** 10.1101/414995

**Authors:** Ronit Dalmat, Negar Makhsous, Gregory Pepper, Amalia Magaret, Keith R. Jerome, Anna Wald, Alexander L. Greninger

## Abstract

HIV drug resistance genotyping is a critical tool in the clinical management of HIV infections. Although resistance genotyping has traditionally been conducted using Sanger sequencing, next-generation sequencing (NGS) is emerging as a powerful tool due to its ability to detect lower frequency alleles. However, the value added from NGS approaches to antiviral resistance testing remains to be demonstrated. We compared the variant detection capacity of NGS versus Sanger sequencing methods for resistance genotyping of 144 drug resistance tests (105 protease-reverse transcriptase tests and 39 integrase tests) submitted to our clinical virology laboratory over a four-month period in 2016 for Sanger-based HIV drug resistance testing. NGS detected all true high frequency drug resistance mutations (>20% frequency) found by Sanger sequencing, with greater accuracy in one instance of a Sanger-detected false positive. Freely available online NGS variant callers Hydra and PASeq were superior to Sanger methods for interpretations of allele linkage and automated variant calling. NGS additionally detected low frequency mutations (1-20% frequency) associated with higher levels of drug resistance in 30/105 (29%) of protease-reverse transcriptase tests and 4/39 (10%) of integrase tests. Clinical follow-up of 69 individuals for a median of 674 days found no difference in rates of virological failure between individuals with and without low frequency mutations, although rates of virological failure were higher for individuals with drug-relevant low frequency mutations. However, all 27 individuals who experienced virological failure reported poor adherence to their drug regimen during preceding follow-up time, and all 19 who subsequently improved their adherence achieved viral suppression at later time points consistent with a lack of clinical resistance. In conclusion, in a population with low antiviral resistance emergence, NGS methods detected numerous instances of minor alleles that did not result in subsequent bona fide virological failure due to antiviral resistance.

**Importance:** Genotypic antiviral resistance testing for HIV is an essential component of the clinical microbiology and virology laboratory. Next-generation sequencing (NGS) has emerged as a powerful tool for the detection of low frequency sequence variants (allele frequencies <20%). Whether detecting these low frequency mutations in HIV contributes to improved patient health, however, remains unclear. We compared NGS to conventional Sanger sequencing for detecting resistance mutations for 144 HIV drug resistance tests submitted to our clinical virology laboratory and detected low frequency mutations in 24% of tests. Over approximately two years of follow-up for 69 patients for which we had access to electronic health records, no patients had virological failure due to antiviral resistance. Instead, virological failure was entirely explained by medication non-adherence. While larger studies are required, we suggest that detection of low frequency variants by NGS presents limited marginal clinical utility when compared to standard of care.

## Introduction

Antiretroviral drug therapy (ART) for HIV has been tremendously successful in reducing HIV morbidity, mortality, and transmission (1). However, rising rates of HIV infections resistant to antiretroviral drugs pose a major threat to ongoing efforts to control the pandemic (2). Routine HIV drug resistance genotyping improves health outcomes in HIV patients (3, 4) and is a cost-effective tool in the clinical management of HIV infection (5).

Genotypic resistance testing assays have traditionally been based on Sanger sequencing (6). While Sanger methods are highly reproducible and validated, minority HIV resistance mutations present in less than 20% of the viral population may escape detection (with a range of approximately 10-30%, depending on the sample context) (6, 7). The capacity of next-generation sequencing (NGS) to provide additional data on low-frequency drug resistant mutations (DRMs) and its potential for lower costs per sample with large batches has led many clinical laboratories to consider transitioning from Sanger sequencing to NGS (8, 9). A commercial NGS HIV drug resistance test has recently received European and Singaporean in vitro diagnostics approval (10). Numerous previous studies have demonstrated NGS methods detect more resistance-associated mutations at low frequencies that Sanger is unable to detect (11-20). However, the predictive value of detectable low frequency mutations for virological failure or other clinical outcomes remains unclear (21-23).

Anticipating the potential utility of NGS for HIV drug resistance detection is also complicated by the ever-changing landscape of HIV therapy. A previous pooled analysis of HIV-1 resistance mutations associated with NNRTI resistance found a dose-dependent increased risk of virological failure with first-line ART (24). However, prior studies seeking to associate low frequency DRMs with virological failure provide insufficient evidence of NGS utility because they lacked an appropriate retrospective comparison group (15), were conducted within a clinical trial (12) (thus lacking representation of real-world patient samples submitted for genotyping in routine care), or were conducted more than 3 years ago and thus focused on NNRTIs (11, 24). Current ART treatment guidelines for treatment-naïve HIV-1-infected patients include an integrase strand transfer inhibitor (INSTI) (25). Dolutegravir is a particularly promising therapy due to its high barrier to resistance (26-28). Given that recent studies of low frequency resistance mutations during integrase and protease inhibitor-based treatments have generally failed to find an association with virological failure (27, 29-34), the clinical value of NGS sequencing for a standard clinical laboratory requires more investigation.

Our reference laboratory provides clinical virology testing for patients in the western Washington region and performed 817 HIV resistance genotyping tests in 2017. Recently, Seattle became one of the first metropolitan areas in the world to attain the 90-90-90 UNAIDS treatment goals, wherein 90% of the patients receiving ART achieve viral suppression (35). Clearly, these goals were reached in the absence of NGS drug resistance testing. Given the concomitant interest in the perceived superiority of NGS for drug resistance in the setting of already successful drug resistance management, we sought 1) to evaluate the technical capacity of Sanger versus NGS sequencing methods for samples submitted to our clinical virology laboratory for HIV resistance genotyping, and 2) to assess the clinical impact of low frequency DRMs detected only by NGS on viral suppression over time.

## Materials and Methods

### Test cohort

Serum/plasma for HIV antiviral resistance testing were collected as part of routine care and Sanger sequenced for antiviral resistance in our reference clinical virology laboratory using a nested RT-PCR approach described below (Figure 1A)(36). To best compare Sanger versus NGS and to limit PCR cycles prior to NGS library preparation for more accurate allelic representation, we used stored first-round RT-PCR amplicons from prior HIV drug resistance testing: 105 protease-reverse transcriptase (Pr-RT) and 39 integrase (INT) amplicons. To allow clinical follow-up while maintaining drug resistance testing relevance for current HIV regimens, we performed NGS testing on amplicons from tests originally performed approximately two years ago (February to May 2016). Ethical approval for this study was granted by the University of Washington Institutional Review Board.

**Figure 1.**
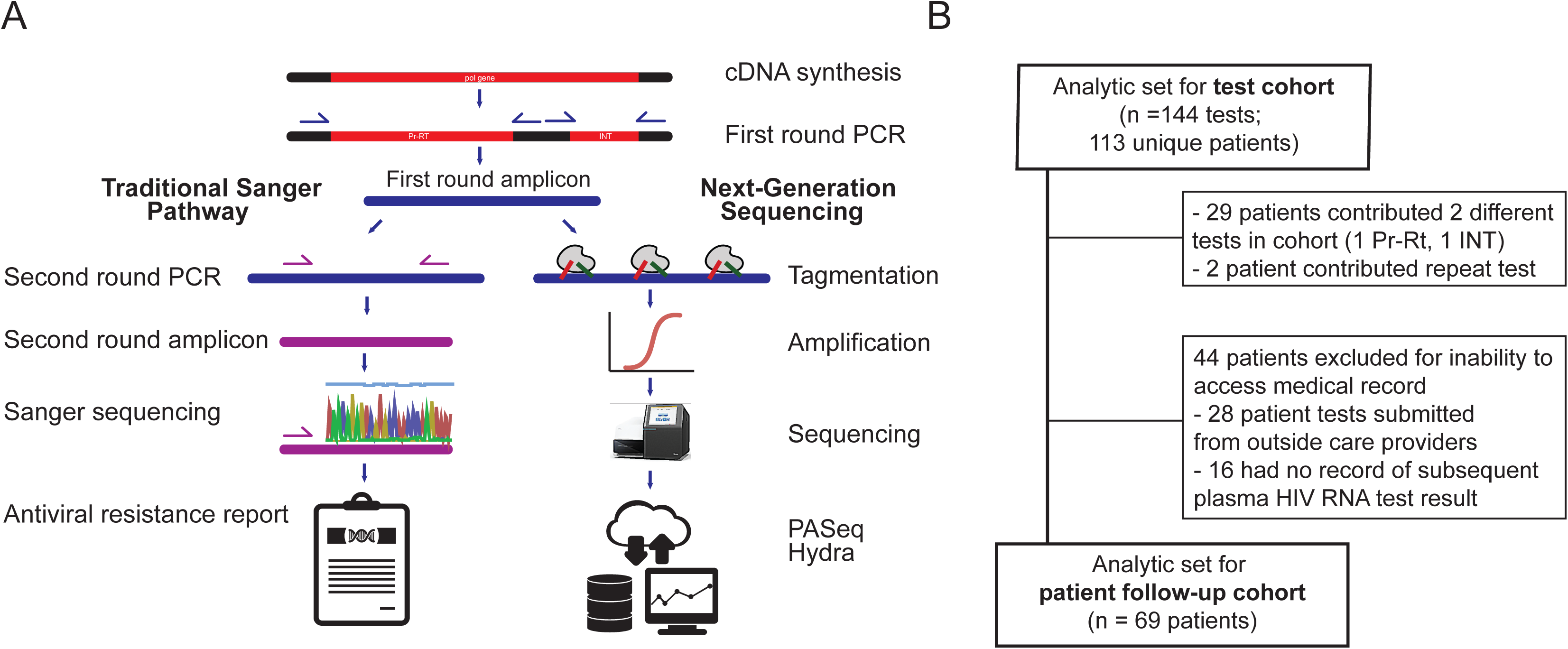
Sequencing protocol comparison. A) First-round amplicons from clinical resistance genotyping by Sanger sequencing were also processed by next-generation sequencing and the resultant resistance profiles compared. B) Test cohort included drug resistance genotyping tests originally performed as part of routine HIV care at UW Medicine between February and May 2016, and patient cohort included those patients who provided tests in the test cohort and had available follow-up records between sample date and May 25, 2018.

### Sanger sequencing, variant calling, and interpretation

Results of previous genotypic resistance assays performed by our clinical virology laboratory were obtained as the standard of care comparison group. Plasma/serum were previously processed through the laboratory’s standardized HIV genotypic resistance assay protocol, which involves HIV RNA separated from plasma via the Boom method silica extraction (37). Extracted RNA was used in a RT-PCR reaction using random hexamers to create cDNA with reagents from GeneAmp RNA PCR kit (Perkin-Elmer, Foster City, CA) according to the manufacturer’s protocol.

Nested PCR amplification was performed in 50ul volume reactions, consisting of 5ul 10X PCR buffer (with 15mM MgCl_2_), 8 ul dNTP’s (1.25mM each), 1ul each forward and reverse primer (20pmol/ul), 0.5 ul Taq (2.5 units per 10μl cDNA sample for first-round PCR; 2.5 units per 5μl PCR product for second round PCR), and nuclease free water. A list of primers is provided in Supplementary Table S-1. A thermal cycler protocol programmed for 94°C x 5 minutes for initial denaturation, followed by 35 cycles of 15 seconds at 94°C for denaturation, 30 seconds at 57°C for annealing, 2 min at 72 °C for extension, and final extension at 72 °C for 7 minutes. An annealing temperature of 55 °C was used for second round nested PCR.

A sample of the first-round PCR product was stored in a −20°C sample archive. The second-round PCR product was visualized on a UV light box (254nm wavelength) after gel electrophoresis [1.5 % agarose ethidium bromide gel with 1X Tris Acetate-EDTA buffer (TAE) (Sigma T9650-1L) run at 80mV for 30 minutes. This PCR product was cleaned by adding 2 ul of Exonuclease I and Shrimp Alkaline Phosphatase directly to 10 ul PCR product without changing buffer conditions. Thermal cycler programmed for 37°C x 15 minutes, 80°C x 15 minutes, 4° C x infinity. Each sample was diluted with nuclease free water to achieve a final concentration 10-40 ng per reaction. Each sequencing primer (5.0 pmol/uL) was added to 10ul of the pre-cleaned diluted PCR template in a pre-labelled PCR Genemate Skirted 96 well PCR plate (T-3107-1).

The samples were sent to a centralized sequencing facility. A Big Dye terminator kit was used for the sequencing reaction. Montage SEQ96 Millipore sequence clean up kit was used for post sequencing. Samples were analyzed using 3730 XI ABI instrument and sequence analysis performed using Sequencher software v5.2.2

Criteria for acceptable data included: raw data peak intensities between 1000 and 5000, average noise intensity <15%, pure base QV rated high and no abrupt signal changes, elevated base line or spikes. Variant mixes were called when visible in both forward and reverse reads, with and minor peaks visible for at least 25% surface area of the major peak.

Results were recorded in a genotypic resistance assay report that was returned to the patient’s care provider via the electronic medical record. The report includes a list of mutations associated with drug resistance in the three *pol* gene segments (protease, reverse transcriptase, and integrase) and an interpretation of the resistance profile (what drugs should be avoided) based on the detected mutations. Mutations listed in the genotypic resistance assay report were also interpreted using the Stanford HIV Drug Resistance Database (HIVdb version 8.5) (38). For each mutation, levels of resistance were recorded categorically according to the HIVdb standard categories: (0) no evidence of resistance; (1) potential low-level resistance; (2) low level resistance; (3) intermediate resistance; and (4) high level resistance. A sample was considered to have any resistance if it had one or more mutations categorized as a level 1 or higher.

### NGS sequencing, variant calling, and interpretation

First-round amplicons for each test were retrieved from −20°C archive and cleaned using 1.0X Ampure beads, quantitated on a Qubit 3.0, and diluted to 1 ng/uL. Libraries were prepped using quarter-reactions of Nextera XT followed by 15 cycles of dual-indexed PCR amplification and sequenced to achieve between 50,000-100,000 reads per sample on an Illumina MiSeq using both 1×192 and 2×300 bp runs. Amplicons with less than 10,000x coverage were excluded to ensure sufficient coverage for detection of low frequency mutations. Samples were sequenced in batches of 20-24 samples to minimize the possibility of index cross-talk on the MiSeq platform and each included positive (8E5 cell line) and negative (nuclease-free water) controls to confirm the run. Sequences were preprocessed using cutadapt (39) and uploaded to two online variant callers: PASeq (https://www.paseq.org/) and HyDRA (https://hydra.canada.ca/). These two variant callers were selected because they are highly-developed, free variant callers with user-friendly web interfaces that require minimal bioinformatics skills and provide robust, reproducible, and easy-to-interpret results that could be implemented in a clinical laboratory (40). Both callers use an annotated HXB2 sequence as their reference for variant calling and the well-established Stanford HIV Drug Resistance Database (version at time of use: 8.5) to provide resistance interpretations for DRMs.

A “low frequency” DRM was defined as a resistance-associated mutation between 1% and 20% allele frequency, based on the default 1% minimum allele frequency needed for a mutation to be considered in the drug resistance report produced by both HyDRA and PASeq (40, 41). A DRM was considered “high frequency” if detected at greater than 20% allele frequency. Resistance interpretations were determined by the same method described above for Sanger-detected mutations.

### Patient characteristics and clinical record abstraction for patient follow-up cohort

Patients’ age, sex, and sample date (baseline) were determined based on patient demographics recorded on the genotypic resistance assay report for all patients included in the test cohort. Electronic medical records were reviewed for clinical outcomes. Patients lacking a recorded plasma HIV RNA test result between the sample date and May 25, 2018 were excluded from the patient follow-up cohort due to lack of available follow-up. Laboratory and clinical visit records for patients included in the follow-up cohort were abstracted using a structured data form (information collected is described below), which was designed based on a preliminary in-depth search of five records prior to full record search and abstraction.

The following information was recorded for each patient: date of first plasma HIV RNA test in our laboratory information system, plasma HIV RNA load at date of sample (or nearest date), drug regimen prescribed by clinician during clinic visit subsequent to sample date (when both plasma HIV RNA measurement and HIV genotyping results were available to inform clinician actions), and date of most recent office visit where HIV care was received. Adherence (yes/no) was assessed based on clinician visit notes as to whether the patient reported being adherent to the drug regimen prescribed at each clinic visit after baseline. Treatment experience was assessed as (naïve/experienced) based on whether a new diagnosis or initiation of first treatment was noted in the clinical record at baseline clinic date.

The primary outcome of interest, virological failure (yes/no), was determined according to the U.S. Health and Human Services definition—failing to achieve or maintain suppression of viral replication to a plasma HIV RNA level <u>≤</u>200 copies/mL (42)—at a test date more than one month after beginning treatment regimen. Prescribed drug regimens were extracted from the medical record. Low frequency mutation exposure was categorized as present/absent, according to whether the Stanford HIVdb resistance profile determined by NGS was higher in level of resistance compared to the profile reported by Sanger data in the clinical record. In a second calculation of risk, exposure was determined as present when a patient was prescribed a drug regimen which would have been contraindicated based on the resistance profile (including low-frequency mutations) found by NGS.

### Statistical analyses

Agreement of DRMs called by each NGS caller (HyDRA and PASeq) was analyzed for concordance using percent agreement and Cohen’s Kappa (a standard agreement coefficient for binary ratings). The PASeq and HyDRA allele frequency measures for all called mutations were compared by linear regression analysis. Any variants not called by either caller were assumed to be 0% allele frequency and excluded from the concordance analysis due to absence of quantitation. The linear model coefficient (HyDRA’s allele frequency measurement as a function of PASeq’s) was also reported. Incidence rate of virological failure was determined based on patients’ first occurrence of HIV RNA level ≥200 copies/mL divided by time at risk (between sample date and censored date). Patients were censored at date of measured virological failure or last available plasma HIV RNA quantification test prior to May 25, 2018. Relative virological failure rate ratios were calculated using the *fmsb* package (43) and all statistical analyses were conducted in R (version 3.4.3) through the RStudio interface (version 1.0.153).

## Results

### NGS estimated higher levels of drug resistance in test cohort

Of 144 total tests, 29 patients had both Pr-RT and INT tests ordered for them during the sample period and two patients had a repeat Pr-RT test ordered in the sample period. Therefore, a total of 113 unique patients were ultimately included in the test cohort (Figure 1B). Patients were predominantly male (91%) with a median age of 40 (range, 5-67). Most (95%) had HIV subtype B, 5 patients had subtype C (contributing 6 tests), and one had subtype A (1 test). Additional descriptive statistics for the test cohort are listed in Table 1.

**Table 1.** Demographic and clinical characteristics of patients included in the test cohort.

| Patient characteristics<sup>1</sup> | (n = 113) |  |
| --- | --- | --- |
| Patient age, median (IQR) | 40 (33-49) |  |
| Men, n (%)<sup>1</sup> | 103 (91) |  |
| Plasma HIV RNA at baseline [log10(copies/mL)] – median (IQR) | 4.2 (3.7-4.8) |  |
| Missing plasma HIV RNA measure at baseline | 27 (24) |  |
| B subtype, n (%) | 107 (95) |  |
| C subtype | 5 |  |
| A subtype | 1 |  |
| Test characteristics | (n = 144) |  |
| Any (≥1) DRM reported by clinical genotyping (Sanger sequencing), n (% of tests) | 40 (28) |  |
| PI | 2 (2) |  |
| NRTI | 18 (17) |  |
| NNRTI | 28 (27) |  |
| INSTI | 1 (3) |  |
|  | HyDRA<sup>2</sup> | PASeq |
| Any (≥1) DRM detected by NGS, n (% of tests) | 60 (42) | 63 (44) |
| PI | 11 (11) | 11 (10) |
| NRTI | 27 (26) | 24 (23) |
| NNRTI | 36 (34) | 39 (37) |
| INSTI | 5 (13) | 4 (10) |

<sup>1</sup> Percentages calculated based on 113 unique patients in test cohort: 29 patients had both
Pr-RT and INT tests and 2 patients had a repeat Pr-RT test during the sample period.
(*Protease-reverse transcriptase test*) PI: protease inhibitors; NRTI:
nucleoside/nucleotide reverse transcriptase inhibitors; NNRTI: non-nucleoside
reverse transcriptase inhibitors
(*Integrase test*) INSTI: integrase inhibitors
<sup>2</sup> HyDRA DRMs were used as the “NGS” value for subsequent analyses in the patient follow- up cohort that compared Sanger to NGS resistance profiles.

Using the standard of care genotyping test by Sanger sequencing, 40 (28%) tests were positive for resistance to at least one drug. By NGS, 63 (44%) tests were positive for resistance by PASeq and 60 (42%) by HyDRA. NNRTI resistance was most common among both methods; PI and INSTI resistance were least common (Table 1). Of 105 Pr-RT tests, Sanger detected 2 (2%) with PI DRMs, 18 (17%) with NRTI DRMs, and 28 (27%) with NNRTI DRMs. Among 39 INT tests, 1 (3%) had a DRM associated with resistance to INSTI. PASeq detected 9 (9%) more samples with PI resistance; 6 (6%) more samples with NRTI resistance, 11 (10%) more samples with NNRTI resistance, and 3 (8%) more samples with INSTI resistance. HyDRA found similarly higher resistance compared to Sanger: 9 (9%) more samples with PI DRMs, 8 (8%) more with NNRTI DRMs, 7 (9%) more with NRTI DRMs, and 4 more (10%) with INSTI DRMs. Overall, NGS estimated higher levels of drug resistance to one or more antiretroviral drugs for 34 (24%) tests—30/105 (29%) protease-reverse transcriptase tests and 4/39 (10%) of integrase tests.

### Sanger and NGS variant calls have high concordance with some limitations

Sanger and NGS showed almost perfect agreement in their detection of high frequency DRMs (97% of tests). In total, four unexpected differences were observed in a comparison of Sanger versus NGS-detected variant calls (overview in Figure 2 and Figure S-1; detailed in Figure 3A-D). All were the result of position-specific over-interpretation of variants in Sanger. One Pr-RT test was discordant in the mutation called, resulting in a false-positive error by Sanger for antiviral susceptibility. Sanger called a mixed L210CW mutation, which is associated with high resistance to didanosine (ddI), abacavir (ABC) and tenofovir (TDF) (38). NGS, however, did not call this mutation (Figure 3A). Upon manual inspection of the NGS sequence, it became clear that the Sanger read was a false positive for resistance due to the inability of Sanger to interpret linkage between two adjacent nucleotide bases. In this instance, the reference codon was TTG (L - leucine) but the viral population contained a mix of the reference codon and TGT (C - cysteine). Thus, NGS read the variant as L210C and did not call it as a DRM. Sanger, on the other hand, interpreted the codon as having a mix of Gs at both the second and third bases, which led to an interpretation of TGT (C - cysteine) and TGG (W – tryptophan). In three other Pr-RT tests, Sanger reported mutations that were detected at less than 20% allele frequency by NGS: V108I was called by HyDRA (15.9%) and PASeq (17.1%) (Figure 3B), K103N was called by HyDRA (8.9%) and PASeq (7.0%), and H221Y was called by HyDRA (17.9%) and PASeq (17.7%) (Figure 3C). These results demonstrate that DRMs at allele frequencies under 20% can be detected by Sanger sequencing.

**Figure 2.**
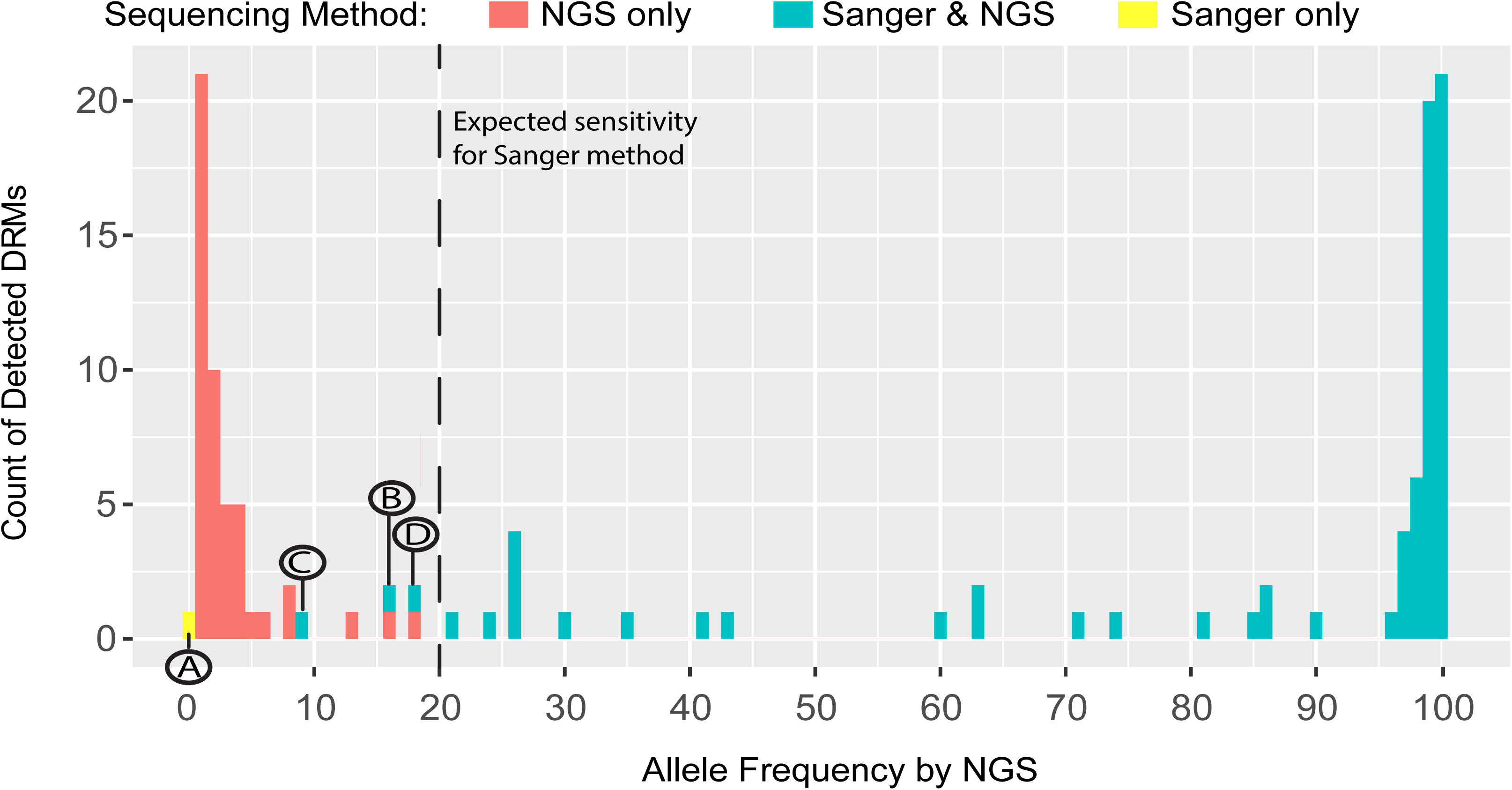
Histogram of DRM frequency by method of sequencing. Number of DRMs detected at each allele frequency, by NGS only (red) by both Sanger and NGS (blue), and by Sanger only (yellow).. A-D letters highlight disagreements found between NGS and Sanger variant calls made at the 20% allele frequency threshold expected for Sanger sequencing. Each instance is detailed in the corresponding letter of Figure 3.

**Figure 3.**
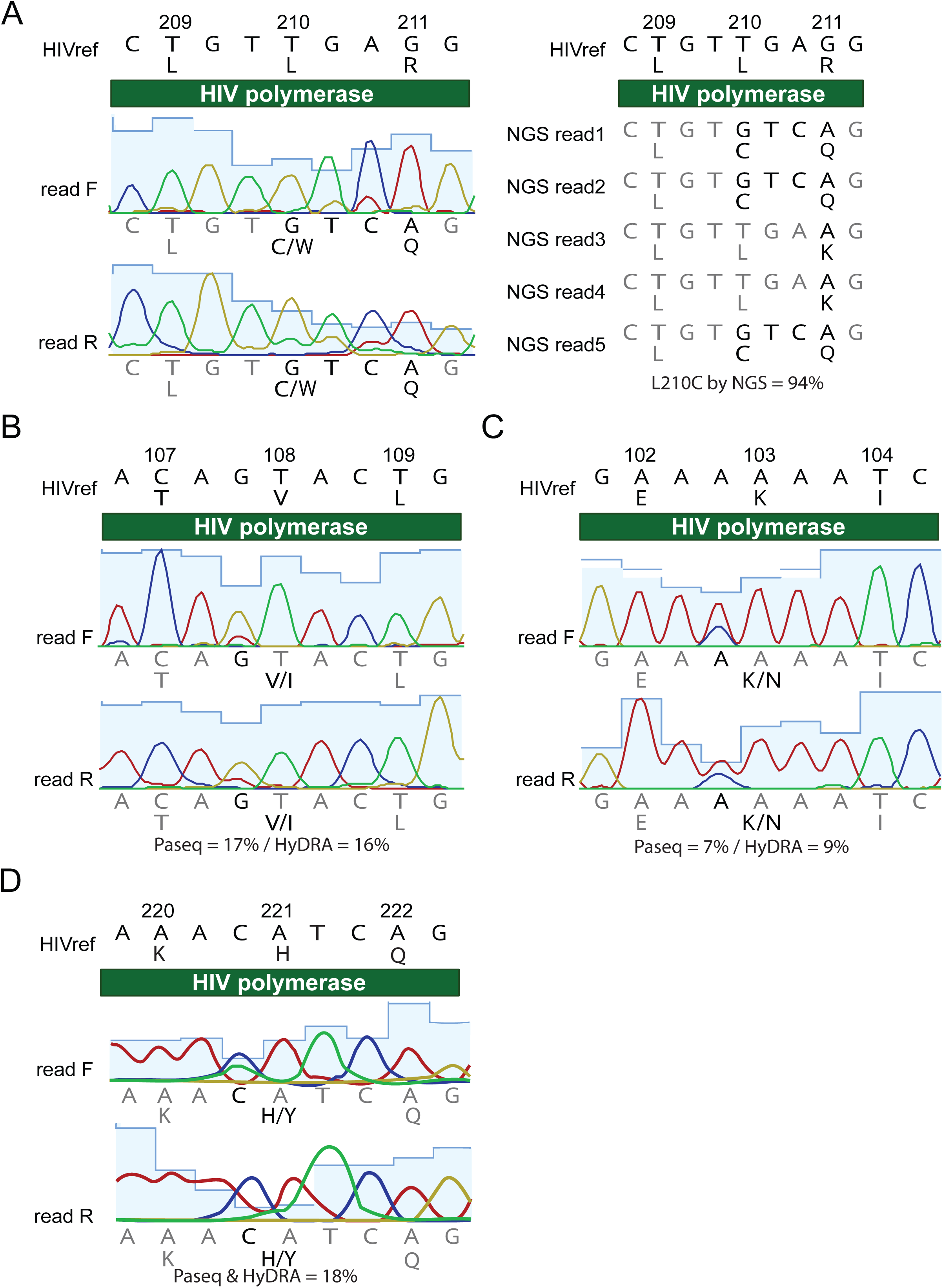
Discrepancies between Sanger and NGS variant calls at 20% allele frequency threshold. A) Sanger called a mixed L210CW variant, which is associated with resistance to three NRTI drugs. However, neither NGS caller called the variant as a DRM because they recognized the linkage between two adjacent nucleotide bases, which clarified the variant as TGT (C – cysteine; polymorphism), rather than a mixture that included TGG (W – tryptophan; DRM). Three different DRMs were called by Sanger sequencing that were detected at less than 20% allele frequency by NGS, including V108I (16-17%, B), K103N (7-9%, C), and H221Y (18%, D).

### NGS callers HyDRA and PASeq are essentially equivalent in their DRM calls

At a 20% allele frequency threshold, HyDRA and PASeq had 100% agreement in terms of which mutations they detected (binary: detected/not; Kappa = 1.0). Concordance of DRM allele frequencies (continuous: 0-100%; Bland-Altman plot in Figure 4A) detected by HyDRA versus PASeq was also very high (linear model coefficient = 1.00; R^2^ = 1.00). Of note, HyDRA reported many more accessory variants and polymorphisms compared to PASeq (Figure 4B). Of 56 unique DRMs detected among 105 Pr-RT sequences and 3 unique DRMs detected in 39 INT sequences, 40 (71%) and 3 (100%) mutations, respectively, were detected with perfect agreement between the callers (Kappa = 1.0).

**Figure 4.**
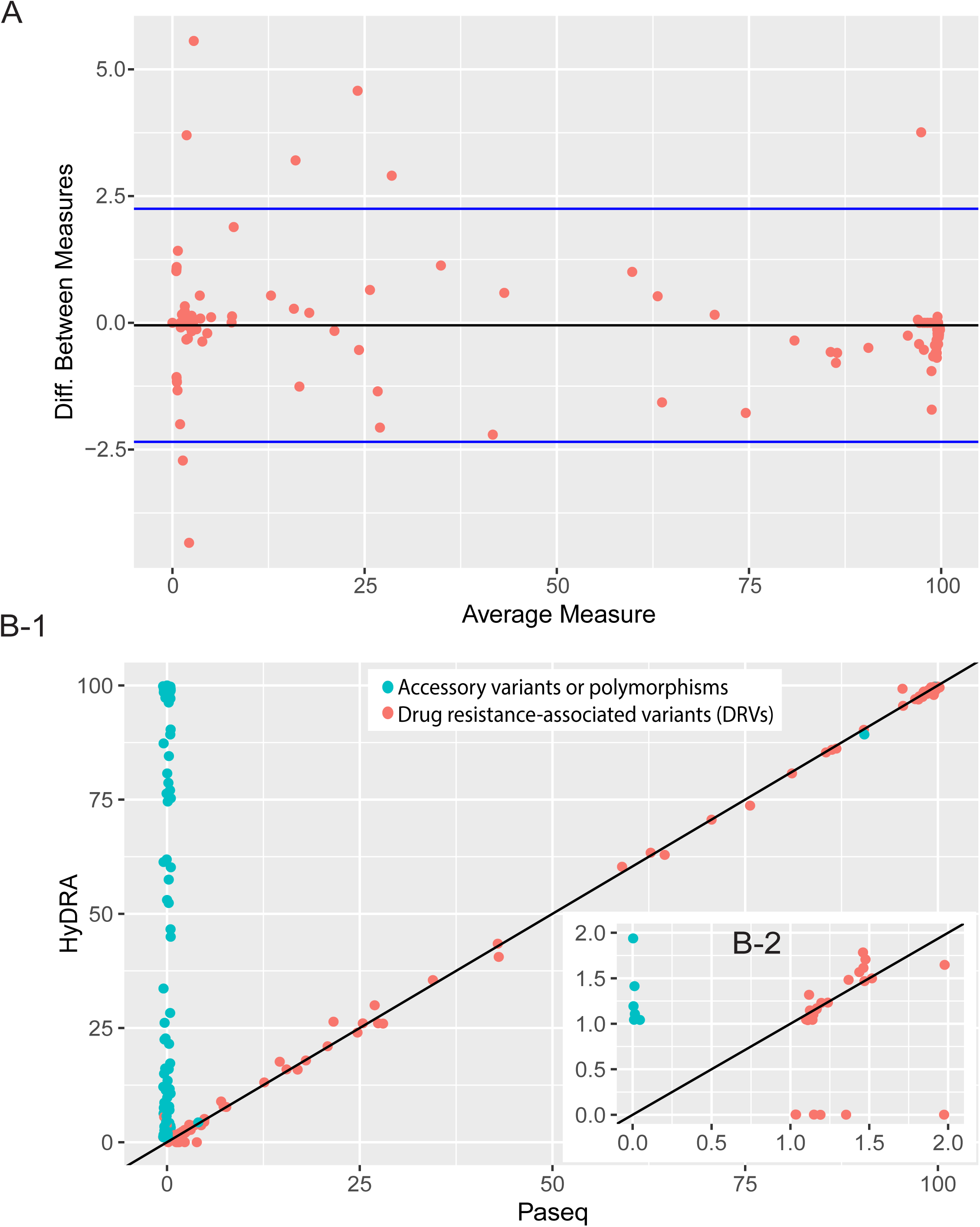
Concordance of allele frequency calls by HyDRA and PASeq variant callers. A).Bland-Altman plot comparing allele frequency measurements for variants called by PASeq and HyDRA. B-1) Correlation plot of allele frequency measurements for variants called by PASeq (x-axis) versus HyDRA (y-axis). Accessory mutations are in blue and DRMs are labeled in red. The default setting of HyDRA includes calls of many more accessory variants than PASeq, which led to the large number of accessory mutations called by HyDRA but not by PASeq. The accessory mutation S168G (blue) was called by both callers at 90% allele frequency. B-2) Zoomed in view of 1-2% frequency range (cluster of observations in bottom left corner of Figure 4B-1). *For figure illustration, all true zero values were assigned a random uniform value between 0-1% in (A) and 0-0.1% in (B).

DRMs on which the two callers disagreed were predominantly very low frequency calls (Figure 4B); the median frequency of mutations detected only by either HyDRA or PASeq (and thus “missed” by the other) was 1.15%. Upon manual inspection of DRMs between 1-2% allele frequency, the discrepancies were attributable to minute differences in the number of reads registered by each caller. For further comparisons of NGS and Sanger results, the callers were assumed to have performed sufficiently similarly and thus the variant and resistance profiles called by HyDRA were used to represent the NGS results for subsequent analyses comparing NGS to Sanger in the patient follow-up cohort.

### Patient follow-up cohort included 69 patients followed for an average of 674 days

Of 113 patients included in the test cohort, 69 received care at UW Medicine and had obtainable clinical follow-up data. The median follow-up time was 674 days (IQR: 560-728 days). This cohort was a subset of the test cohort, with a similar age and sex distribution (median age = 40; 88% men) and prevalence of low frequency DRMs (32% of patients). Summary statistics for the cohort are provided in Table 2. A higher proportion of patients with low frequency DRMs were female, compared to those without. In the patient cohort, 27 patients (39%) experienced virological failure of any plasma HIV RNA ≥200 copies/mL during the follow-up period, 12 (17%) patients had changes to their drug regimen in follow-up and 12 (17%) patients received additional Sanger resistance tests as part of regular care. None of these additional Sanger resistance tests detected any additional DRMs compared to the baseline Sanger resistance test.

**Table 2.**
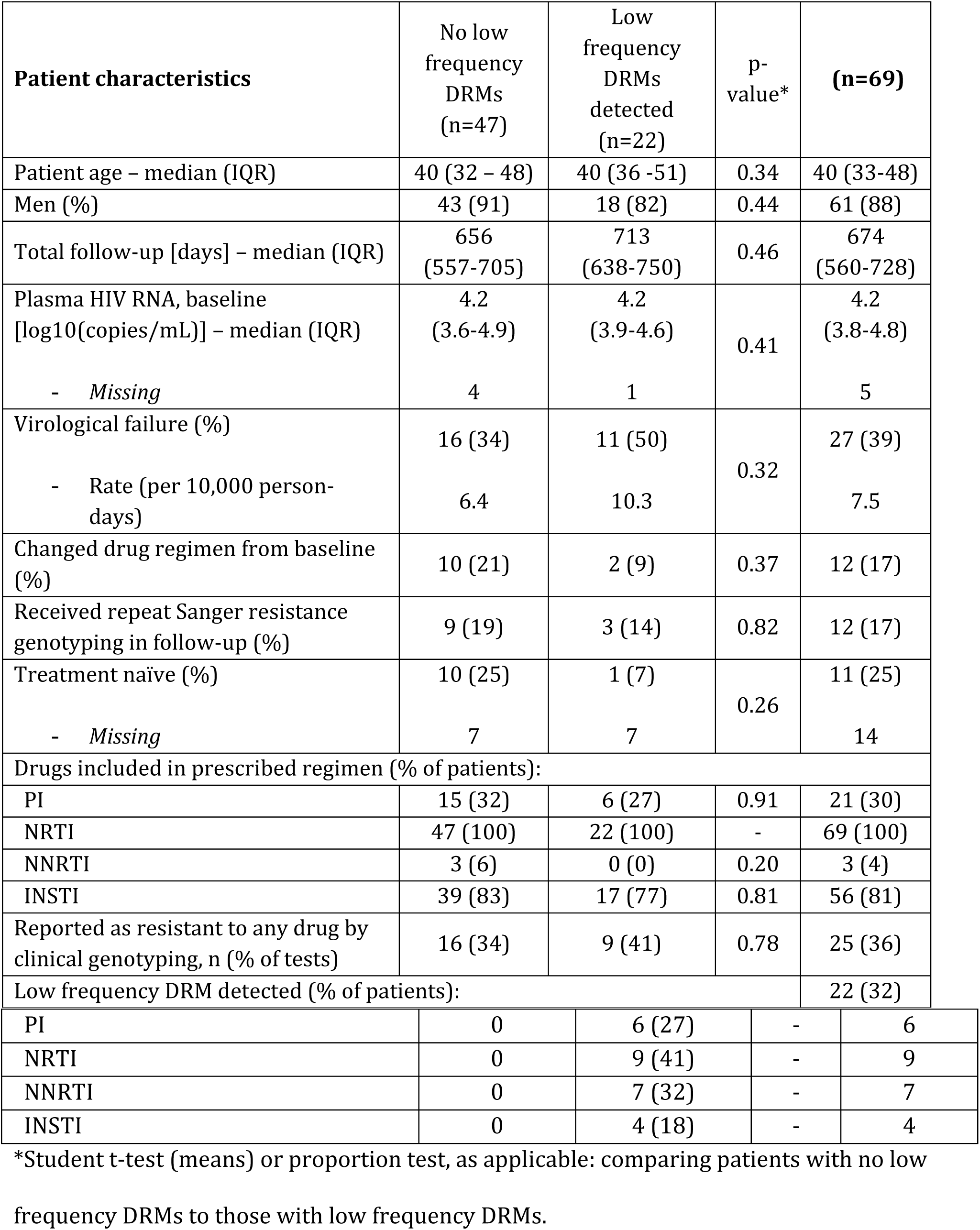
Demographic and clinical characteristics of patients included in the patient follow-up cohort.

| Patient characteristics | No low frequency DRMs (n=47) | Low frequency DRMs detected (n=22) | p-value* | (n=69) |
| --- | --- | --- | --- | --- |
| Patient age – median (IQR) | 40 (32 – 48) | 40 (36 -51) | 0.34 | 40 (33-48) |
| Men (%) | 43 (91) | 18 (82) | 0.44 | 61 (88) |
| Total follow-up [days] – median (IQR) | 656 (557-705) | 713 (638-750) | 0.46 | 674 (560-728) |
| Plasma HIV RNA, baseline [log10(copies/mL)] – median (IQR) | 4.2 (3.6-4.9) | 4.2 (3.9-4.6) | 0.41 | 4.2 (3.8-4.8) |
| - Missing | 4 | 1 |  | 5 |
| Virological failure (%) | 16 (34) | 11 (50) | 0.32 | 27 (39) |
| - Rate (per 10,000 person-days) | 6.4 | 10.3 |  | 7.5 |
| Changed drug regimen from baseline (%) | 10 (21) | 2 (9) | 0.37 | 12 (17) |
| Received repeat Sanger resistance genotyping in follow-up (%) | 9 (19) | 3 (14) | 0.82 | 12 (17) |
| Treatment naïve (%) | 10 (25) | 1 (7) | 0.26 | 11 (25) |
| - Missing | 7 | 7 |  | 14 |
| Drugs included in prescribed regimen (% of patients): |  |  |  |  |
| PI | 15 (32) | 6 (27) | 0.91 | 21 (30) |
| NRTI | 47 (100) | 22 (100) | - | 69 (100) |
| NNRTI | 3 (6) | 0 (0) | 0.20 | 3 (4) |
| INSTI | 39 (83) | 17 (77) | 0.81 | 56 (81) |
| Reported as resistant to any drug by clinical genotyping, n (% of tests) | 16 (34) | 9 (41) | 0.78 | 25 (36) |
| Low frequency DRM detected (% of patients): |  |  |  | 22 (32) |
| PI | 0 | 6 (27) | - | 6 |
| NRTI | 0 | 9 (41) | - | 9 |
| NNRTI | 0 | 7 (32) | - | 7 |
| INSTI | 0 | 4 (18) | - | 4 |

\*Student t-test (means) or proportion test, as applicable: comparing patients with no low frequency DRMs to those with low frequency DRMs.

### NGS detected more resistance mutations but these did not significantly associate with virological failure, which was more likely attributable to drug non-adherence

Of 69 patients with clinical follow-up, 22 (32%) had low frequency DRMs detected by NGS that were not found by Sanger. Low frequency DRMs were most commonly associated with NRTI resistance (9 patients), followed by NNRTI (7 patients), PI (6 patients), and INSTI (4 patients). Five (7%) patients received drugs which would have been contraindicated based on the resistance profiles found by NGS.

We were not able to detect a difference in the risk of virologic failure between patients with any low frequency DRMs at baseline compared to those without (RR = 1.61; 95% CI: 0.75-3.48). Three patients had only level 1 low frequency DRMs; when we restrict our exposure definition to patients with level 2 or higher resistance (see definition provided in methods section), the virological failure rate in patients with any low frequency DRMs is not significantly higher than in those without (RR = 1.80; 95% CI: 0.82 – 3.92). Among five patients with low frequency DRMs specifically associated with one of their prescribed drugs, four experienced virological failure. Compared to patients without affected regimens, these five patients had a higher rate of virological failure (RR = 3.61; 95% CI: 1.25-10.44).

However, all participants who experienced virological failure and subsequently reported improved adherence to their drug regimen were able to achieve viral suppression—regardless of whether or not they had low frequency DRMs—even though most of them (16 of 19 patients) did not change drug regimens in that time period (Figure 5A).

**Figure 5.**
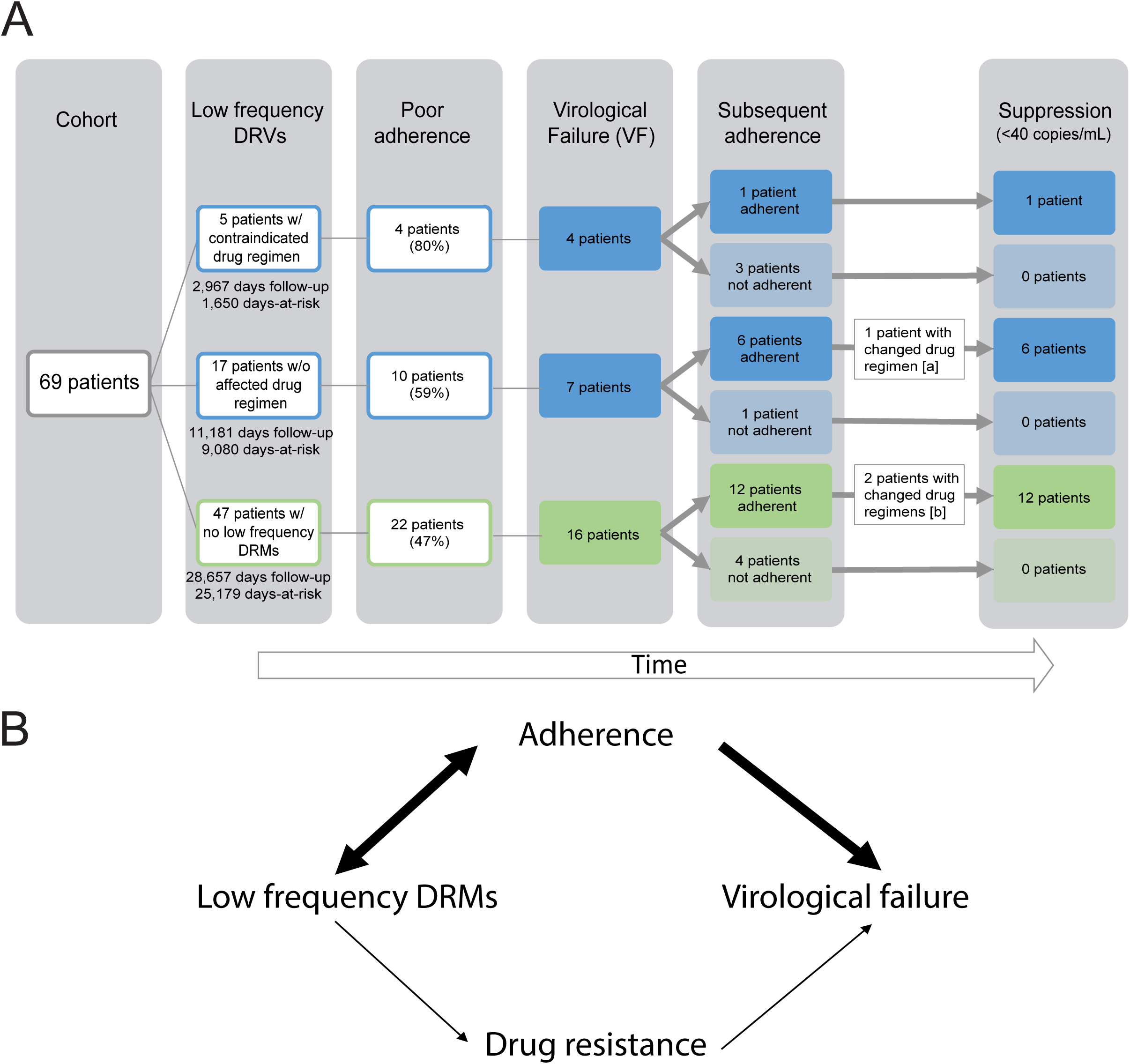
Results of clinical follow-up for patients and theoretical model. A) Low frequency DRMs are correlated with ongoing medication adherence, as a result of prior drug experience and current selective drug pressure, which is thus associated with virological failure, regardless of drug resistance. B) Stratification of clinical outcomes in patient cohort, given the presence or absence of low frequency DRMs associated with reduced susceptibility to their drug regimen. Adherence was defined as documentation of poor medication adherence in clinical chart. Virological failure was defined as plasma HIV RNA level >200 copies/mL at a test date more than one month after baseline. [a] Lamivudine/zidothymidine was replaced with emtricitabine/tenofovir following poor adherence in patient’s regimen, potentially contributing to subsequent suppression. [b] Darunavir was replaced with dolutegravir in both patients’ regimens; potentially contributed to subsequent suppression Days-at-risk refers to time between sample date and censor date. Patients were censored at date of measured virological failure or last available plasma HIV RNA quantification test. Days follow-up includes all days between sample date and last plasma HIV RNA quantification test date.

## Discussion

Our study suggests a limited marginal utility of next-generation sequencing compared to Sanger sequencing for determining HIV antiviral drug resistance in a population with a relatively low prevalence of antiviral resistance. We also demonstrate two free, online variant callers PASeq and HyDRA were marginally superior compared to Sanger sequencing for detection of high frequency DRMs. Both NGS callers were highly concordant with each other in respect to their measured allele frequencies for DRMs. Besides HyDRA’s automatic reporting of many more accessory variants and polymorphisms compared to PASeq, the two callers provided nearly interchangeable results.

PASeq and HyDRA improve upon Sanger sequencing’s manual variant calling procedures by being free and automated, and thus faster, easier, and less expensive. Our study also illustrated two limitations of manual examination procedures for interpreting Sanger chromatograms: missing linkage and reliance on trained laboratory professionals. First, Sanger was unable to impute linkage between two alleles, which manifested as a DRM mixture (L210CW) instead of a polymorphism (L210C). In this example case, the erred call was a false-resistance determination that did not negatively impact patient care because the potentially affected drug types are uncommonly prescribed in this care setting, and the erred variant call is associated with only potential and low levels of associated resistance (zidovudine, stavudine, and didanosine) according to HIVdb. Second, Sanger called three resistance mutations that were below Sanger’s expected 20% threshold of sensitivity. This “over-calling” is illustrative of how the procedures rely on a highly-trained lab technician to review DRM regions in a Sanger sequenced resistance test—had the technician not been experienced enough to expect a DRM in that location, it may have been missed. Both manifestations of Sanger “error” resulted from the human element of Sanger variant calling. Given the ease of use and technical capacity of HyDRA and PASeq, the two callers are both pragmatic options for the detection of variants and interpretation of NGS results in clinical laboratory HIV resistance testing.

In our comparison of overall DRM detection rates, NGS detected more mutations associated with higher levels of drug resistance in 30/105 (29%) of Pr-RT sequences and 4/39 (10%) of INT sequences, due to its capacity to detect low frequency DRMs. These findings are consistent with numerous studies in which NGS detected more DRMs compared to Sanger (11-14, 16-20). However, few studies have performed clinical follow up after detection of low frequency DRMs to determine whether the additional DRMs are relevant to patient care or outcomes. Here, we have presented the first study to investigate the effect of low frequency DRMs on viral suppression in patients on multi-drug therapeutic regimens in a pragmatic retrospective cohort of samples submitted for routine HIV antiviral resistance genotyping to a clinical virology lab.

We found that the 7% of patients prescribed a drug regimen to which they may have been contraindicated based on low frequency DRMs detected by NGS may be at higher risk of virological failure compared to patients without drug-regimen associated DRMs. With 69 patients and a 34% rate of virological failure in those *without* DRMs, we had 80% power to detect increases in the virological failure rate among those *with* DRMs at 75% failure rate or higher. All virological failures in the cohort were associated with medication non-adherence. Low frequency DRMs were most commonly associated with NRTI and NNRTI resistance, even though NNRTI drugs were rarely included in drug regimens of patients in the follow-up cohort (3 patients; 4%). This finding is consistent with previous research (20, 44, 45) and suggests that some proportion of DRMs may have originated during prior drug experience or been transmitted upon infection. Low frequency DRMs may also arise de novo as part of natural viral diversification in the body (46). Poor adherence to ART muddles the association between low frequency mutations and virological failure because it can contribute to the emergence of resistant viral subpopulations (observed as low frequency DRMs), as well as be a primary cause of a patient’s high plasma HIV RNA (Figure 5B). The association may also be confounded by higher viral loads, which could increase the likelihood that a minor variant is present in a sufficient quantity for detection by NGS. However, we think the contribution of this source of confounding is minimal in our study because the laboratory acceptance criteria for the test is 1000 copies/mL and just eight (6%) tests were below this threshold. Furthermore, low frequency variants were detected across a range of viral loads (Figure S2).

More than 80% of patients in our cohort were prescribed a regimen that included an integrase inhibitor at baseline. However, despite this high prevalence of INSTI prescriptions and low prevalence of INSTI resistance mutations, 39% of patients in the cohort experienced virological failure (70% of whom were on an INSTI-based regimen), and just four (15%) failures occurred in patients on drug regimens contraindicated by their NGS resistance profile. Furthermore, the majority of patients never changed drug regimens following their “virological failure,” and patients who continued to report poor adherence experienced uncontrolled viral loads well over 200 copies/mL for the duration of follow-up while those with improved adherence achieved suppression (below the limit of detection; <40 copies/mL). This suggests that poor adherence may be a primary factor affecting rates of viral suppression in this cohort, irrespective of the prevalence of low frequency DRMs.

By evaluating the utility of NGS in the context of a pragmatic patient population, we provide a real-world example of its potential to inform clinical care and elucidate factors that could affect its implementation. Our focus on a single high-resource setting is a potential limitation of the study. Seattle is unique in already having achieved UNAIDS goals of 90% of patients on ARV with suppressed virological loads, indicating a smaller potential return on investment for NGS antiviral testing and a low overall prevalence of antiviral resistance. In a high-resource clinical setting, any indication of resistance on a Sanger resistance assay report—even at low levels—is sufficient to rule out that drug for an ARV regimen. DRMs causing any degree of change to the level of resistance interpreted by HIVdb were therefore considered in this analysis. In less resource-rich settings, however, “potential” (level 1) and “low-level” (level 2) resistance may need to be excluded from the analysis because their clinical significance is not well understood and may be overly conservative for prescribing practices in resource-limited settings.

Another limitation of our study is its low power. Despite significant efforts exerted to obtain a sufficiently large cohort size for the analysis, more than a third of patients in the test cohort were excluded from the sample cohort due to lack of follow-up (or inability to obtain medical records). The exclusion of these individuals is not a major concern as a source of selection bias because loss to follow-up is a regular occurrence in the course of clinical management and remains representative of a pragmatic population receiving testing in a clinical lab. Given the clinical and potential economic importance of this study’s objectives, a long-term cohort study involving many more patients receiving HIV care and repeat sequencing is needed. However, we also caution that such a study may have limited generalizability in the current age of fast-changing HIV clinical pharmacology. As has been the case for efforts to interpret prior similar studies conducted in populations and time periods without integrase inhibitors (11, 24), evolving treatment regimens make it difficult to develop informative pragmatic cohort studies to validate the clinical relevance of low frequency variants.

Regardless, our findings demonstrate two online, free, and easy-to-use NGS variant callers have high concordance with Sanger in a pragmatic clinical setting and thus may be good candidates for implementation as part of a clinical laboratory analysis pipeline for HIV drug resistance. However, given growing prevalence of integrase inhibitor regimens with high barriers to resistance and high prevalence of low frequency DRMs (predominantly NNRTI and NRTI) in patients with poor adherence, we suggest clinical laboratories use caution in overvaluing the low-frequency variant detection afforded by NGS for clinical antiretroviral resistance genotyping as it may not provide much improvement over standardized Sanger methods for detection of clinically relevant resistance in a real-world context.

## Acknowledgements

The authors would like to thank Ryan C. Shean for help with data and metadata submission to NCBI as well as the patients and families served by the University of Washington Virology Lab. ALG reports prior consulting for Abbott Molecular.

## Supplemental Material

**Figure S-1**. **Agreement of Sanger and NGS variant resistance interpretations at 20% and 1% cutoffs.** Low frequency DRMs refer to those between 1-20% frequency and were only called by NGS. Tests with agreement between Sanger and NGS (green); disagreement (blue); Sanger false-positive due to inability to detect linkage (orange).

**Figure S-2. Low frequency variants detected in patients with a range of viral loads.** A) Boxplot distribution of patients’ viral load [plasma HIV RNA log(copies/mL)] at sample date for samples with and without detected low frequency DRMs. B) Viral load of patient in which a low frequency DRM was detected and the allele frequency of the detected DRM(s).

